# Presenilin 1 (PS1) located at mitochondrial inner membrane regulates mitochondrial cristae junction proteins arrangement and cristae formation in HEK293 cells

**DOI:** 10.64898/2026.03.05.709976

**Authors:** Pan You, Pengjin Zhu, Hui Yu, Luwen Wang, Bo Su

## Abstract

Presenilin 1 (PS1), a key pathogenic factor in familial Alzheimer’s disease, is implicated in regulation of mitochondrial functions, yet its precise sub-mitochondrial localization and underlying mechanisms remain poorly understood. In this study, we generated PS1 knockout (PS1 KO) cell lines to investigate the role of PS1 in mitochondrial structure and function. Our results demonstrated that PS1 is directly localized to the mitochondrial inner membrane. PS1 deficiency led to reduced ATP production, impaired mitochondrial respiration capacity, decreased mitochondrial membrane potential, disrupted Ca^2+^ homeostasis, and elevated reactive oxygen species (ROS) accumulation. Moreover, loss of PS1 caused abnormal mitochondrial cristae structure. Further analysis revealed that PS1 interacts with mitochondrial inner membrane proteins. Its absence promotes ATAD3A oligomerization and disrupts its arrangement at mitochondrial cristae junctions, leading to expansion of the mitochondria-associated membrane (MAM) and instability of mitochondrial DNA (mtDNA). Our findings demonstrate that PS1 acts as a central regulator of mitochondrial cristae morphogenesis by modulating protein interaction networks at cristae junctions, thereby illuminating fundamental molecular mechanisms contributing to mitochondrial dysfunctions in Alzheimer’s disease.

## Introduction

Alzheimer’s disease (AD), the predominant form of dementia, manifests as progressive deterioration of cognitive and mnemonic functions[1]. While most cases occur sporadically, approximately 1-5% of early-onset AD arises from autosomal dominant mutations, predominantly in the PSEN1 gene encoding presenilin 1 (PS1)[2, 3]. As a multi-pass transmembrane protein, PS1 constitutes the catalytic core of the γ-secretase complex, which executes the terminal cleavage of amyloid precursor protein (APP) to generate amyloid-β (Aβ) peptides[4, 5]. Pathological accumulation of neurotoxic Aβ species, culminating in amyloid plaque deposition, represents a hallmark neuropathological feature of AD[1, 2]. Beyond its canonical role in amyloid-β production, PS1 regulates multiple cellular processes. It has been particularly investigated that PS1 is involved in modulation of calcium (Ca²□) homeostasis, PS1 could interact with inositol trisphosphate receptor (IP3R)[6], and SERCA2b[7], functioning as a passive calcium channel on the endoplasmic reticulum (ER). AD-associated PS1 mutations perturb ER calcium homeostasis, PS1 dysfunction disrupts mitochondrial-ER Ca^2+^ crosstalk[8] and upregulated function of mitochondrial associated ER membranes[9]. Additionally, PS1 governs lysosomal acidification and calcium dynamics by controlling glycosylation of the v-ATPase V0a1 subunit[10]. Furthermore, emerging evidence indicates direct PS1 localization in mitochondria and mediated mitochondrial functions[11–13]. Wang et al. demonstrated that PS1 could interact with mitochondrial FKBP38 and regulate apoptotic pathways[14]. Recently studies show that pathogenic PS1 variants display detrimental effect on mitochondrial functions[13]. Notably, mitochondrial dysfunction has emerged as a convergent pathological mechanism across neurodegenerative disorders, with impaired bioenergetics, oxidative stress, and aberrant organellar dynamics being consistently observed in AD models and postmortem brains from AD patients[15–17]. Despite decades of investigation into Aβ-centric pathways, the molecular etiology linking PS1 dysregulation to mitochondrial failure remains poorly defined, particularly regarding non-canonical γ-secretase-independent mechanisms.

Mitochondria are double-membrane organelles in which the inner mitochondrial membrane (IMM) forms a highly dynamic and compartmentalized structure essential for energy production, metabolic regulation, and apoptotic control[15, 18, 19]. In neurons, mitochondrial IM integrity is indispensable for sustaining ATP-dependent neurotransmission, calcium ion (Ca²□) homeostasis, and survival pathways critical to these long-lived postmitotic cells[15, 19]. Structural damage to mitochondria is strongly implicated in AD pathogenesis[19]. The IMM achieves functional specialization through cristae invaginations, creating two morphologically and biochemically distinct domains: the boundary membrane (continuous with the outer membrane) and cristae membrane (folded inward), interconnected by cristae junctions. Respiratory chain supercomplexes are enriched in cristae membranes, whereas protein translocases such as TIM complexes predominantly localize to boundary membranes[20, 21]. Cristae junctions, stabilized by the MICOS complex, OPA1, and ATAD3A, are essential for maintaining IMM structural integrity[22, 23]. These protein assemblies cooperatively regulate cristae morphology, directly determining mitochondrial functional efficiency.

ATAD3A is a multi-pass transmembrane protein spanning both mitochondrial membranes[24]. It participates in maintaining inner membrane architecture, stabilizing mitochondrial DNA (mtDNA), and regulating the formation of mitochondria-associated membranes (MAMs) to mediate Ca²□ flux at mitochondria-ER contact sites[25–27]. Furthermore, recent studies demonstrate increased ATAD3A oligomerization in AD patients and models, correlating with MAM expansion and mtDNA instability[28, 29]. ATAD3A further serves as a scaffolding factor critical for the assembly of inner membrane proteins and structural organization of the mitochondrial inner membrane [27]. While ATAD3A is critical for organizing inner membrane protein networks, its essential role in maintaining mitochondrial inner membrane ultrastructure and initiating cristae biogenesis remains elusive.

In this study, we investigated the role of presenilin-1 (PS1) in mitochondrial function and architecture. In HEK293 cells, we found that PS1 localize in mitochondrial inner membrane, and PS1 deficiency induced mitochondrial vacuolization, and impaired mitochondrial function, manifesting as reduced ATP production, disrupted calcium homeostasis, accumulated reactive oxygen species (ROS), and loss of mitochondrial membrane potential. Notably, PS1 deletion increased ATAD3A protein levels and promoted its oligomerization, disrupting the interaction network of mitochondrial inner membrane proteins. These results advance our understanding of PS1-associated mitochondrial dysfunction in AD, underscoring the critical role of PS1 in preserving mitochondrial homeostasis through functional interplay with key regulators such as ATAD3A.

## Materials and Methods

### Plasmids

The PS1 knockout plasmid was constructed by cloning the PS1-gRNA sequence (5’-GCCACGCAGTCCATTCAGGG-3’) into the lentiCRISPR v2 vector, verified by sequencing. The pcDNA3.1 (+)-N4Flag-PS1-wt plasmid was constructed by amplifying the PS1 sequence from cDNA using primers (Forward: 5’-GATCCTCGAGCTACATATTCACCAACCACACCATTGTTGAGGAG-3’, Reverse: 5’-GATCCTCGAGCTAGATATAAAATTGATGGAATGCTAATTGGTC-3’) and cloning it into the BamH I/Xho I-digested vector. The pCMV-N3Myc-MFN2/TIM23 plasmid was constructed by amplifying the MFN2 or TIM23 sequence from cDNA using primers (MFN2-Forword: 5’-GATCAAGCT TATGTCCCTGCTCTTCTCTCGATGCAACTCT-3’, MFN2-Reverse: 5’-GATCCTCGAGCTATCTGCTGGGCTGCAGGTACTGGTGTGT-3’, TIM23-Forword: 5’-GATCAAGCTTATGGAAGGAGGCGGGGGAAGCGGCAACAAAACCAC -3’, TIM23-Reverse: 5’-GATCCTCGAGTCAGAGTGACTGTTGGAGCAAGGAGCCTTTCATGT -3’) and cloning it into the Hind □/Xho I-digested vector. The pFX-mito-EGFP plasmid (Addgene, USA), expressing EGFP targeted to the mitochondrial inner membrane via the COX8 signal peptide, was used for mitochondrial fluorescence labeling. SPLICS Mt-ER Long P2A plasmid (Addgene, USA) was used for mitochondrial associated membrane (MAM) labeling. siRNAs were used for ATAD3A knocking down.

### Cell Culture and Transfection

HEK293 cells were cultured in Dulbecco’s Modified Eagle’s Medium (DMEM) (Macgene, China) supplemented with 10% fetal bovine serum (Lonsera, China) and 1% penicillin-streptomycin (Solarbio, China). The PS1-gRNA-CRISPR-Cas9 plasmid was transfected at a 1:2 ratio using Lipofectamine 2000 (Thermofisher, USA) or PEI MAX (Polyscience, USA). After 48 hours, cells were selected with puromycin (1 μg/mL, Gibco, USA). When cells reached 80%-90% confluence, they were trypsinized, diluted to 0.5 cells/100 μL, and plated at 100 μL per well in 96-well plates. Single-cell clones were expanded and validated for PS1 knockout by Western blot (WB).

### Cell viability assay

Cell viability was assessed using the Cell Counting Kit□8 (CCK□8) assay (Solarbio, China). Cells were seeded in 12□well plates at an identical density and cultured overnight. Then each well was transfected with vector or indicated plasmids under same ratio of PEI. After transfect 6 hours, change the culture medium to a complete culture medium. After transfect 24 hours, cells were incubated with CCK8 dilution (1:10 in culture medium) for 1 hours at 37□°C. To assess background signal, control wells of HEK293 and PS1□KO cells were set up without the addition of CCK□8 reagent. Absorbance was measured at 450 nm using a microplate reader. The absorbance at 450 nm was measured for each well, corrected by subtracting the background absorbance, and normalized to the vector group.

### Dead cell staining

Dead cell was assessed by Hoechst□33342/PI double staining kit (Solarbio, China). Cells were incubated with a staining solution containing Hoechst□33342 and propidium iodide (PI) diluted 1:1000 in culture medium for 30□min at 37□°C under 5% CO□. After washing twice with PBS, cells were maintained in phenol red□free medium (Macgene, China) and imaged with a 10× objective on an IX73 inverted microscope. Dead cells were identified by bright co□staining with both blue (Hoechst□33342) and red (PI) fluorescence.

### Living cell image

For living cell imaging, HEK293 or PS1KO cells was seeded on glass bottom cell culture dishes (cell-NEST, China) and transfected with vector or indicated plasmids. After transfect 48h, cells were used for living cell imaging to measurement mitochondrial functions. For TMRM staining, which was used for mitochondrial membrane potential measurement, cells were incubated with 100 nM TMRM (Thermo Fisher T668, USA) in DMEM medium at 37□, 5% CO_2_ for 10 minutes. As a negative control of TMRM staining, cells were incubated with 20 μM FCCP at 37□, 5% CO_2_ for 30 minutes before TMRM staining. After staining, cells were washed by PBS for three times. Change the medium to a DMEM without phenol red (Macgene, China), all samples were imaged under identical excitation light intensity and acquisition settings by a 60× objective on an IX73 inverted microscope (Olympus, Japan). For mitochondrial Ca^2+^ measurement, Rhod 2 AM dye (AAT Bioquest 21064, USA) was used. cells were incubated with 2 μM at 37□, 5% CO_2_ for 30 minutes. After incubation, washing cell three times with PBS and change the medium to a DMEM without phenol red (Macgene, China), living cell images were captured under identical excitation light intensity and acquisition settings. For mitochondrial reactive oxygen species (ROS) levels measurement, cells were incubated with 1 μM MitoSOX (Thermo Fisher M36008, USA) Red at 37□, 5% CO_2_ for 30 minutes. After incubation, washing cell three times with PBS and change the medium to a DMEM without phenol red, living cell images were captured under identical excitation light intensity and acquisition settings. All images were analyzed via Imagej software under consistent threshold. The fluorescence intensity was averaged to the cell number and normalized to the control group.

### Immunofluorescence

For cell immunofluorescence, HEK293 or PS1KO cell was seed as appropriate density on cell culture plate within glass coverslips, after overnight culture in 37□, 5% CO_2_, cells were transfected at a 1:2 ratio using PEI MAX (Polyscience, USA) and change the culture medium after 6h. After 48h, cells were fixed 30 minutes using 4% PFA (Paraformaldehyde) (ServiceBio, China) at room temperature (RT). Then the cells were permeated using 0.3% Trixton-100 in PBS at RT for 30 minutes. After permeation, cells were blocked by 10% goat serum in PBS (ServiceBio, China) at RT for 1h. After blocking, cells were incubated with anti-body at 4 □ overnight. Washing coverslips with PBS for three times each time 5 minutes at RT. Incubate fluorescent coupled secondary antibody for 2.5 hours at 37 ℃. After washing with PBS three times for 5 minutes each at room temperature, the coverslips were mounted with DAPI-containing mounting medium (Abcam, UK). The immunofluorescent picture was taken by LSM900 laser confocal microscope (Zeiss, Germany). For super-resolution capture, choose Airyscan 2 model with resolution option. Any measurement of fluorescent picture was taken by Imagej software. Mitochondrial morphology was analyzed using the MitochondriaAnalyzer v2.3.1 plugin with default threshold settings[30]. Mitochondria that were completely fragmented and exhibited perinuclear clustering were defined as punctation. MAM area was quantified in ImageJ using uniform thresholding.

### Mitochondrial Isolation

The mitochondria were isolated from cells as described previously[31]. For cell mitochondria isolation, 90 dishes of PS1KO cells were transfected with Flag-PS1-wt plasmid at a 1:2 ratio using PEI. After 48 hours, cells were harvested, resuspended in isolation buffer (225 mM mannitol, 75 mM sucrose, 0.1 mM EGTA, 30 mM Tris–HCl, pH 7.4), and homogenized with a Teflon glass homogenizer. The homogenate was centrifuged at 600 × g for 5 minutes at 4 °C twice to collect the supernatant. The supernatant was then centrifuged at 7000 × g for 10 minutes. The pellet (crude mitochondria) was washed twice with starting buffer (225 mM mannitol, 75 mM sucrose, 30 mM Tris–HCl, pH 7.4) and resuspended in MRB (250 mM mannitol, 5 mM HEPES, pH 7.4, 0.5 mM EGTA). The supernatant containing ER and cytosolic fractions. To perform such sub-fractionation centrifuge supernatant at 20,000 × g for 30 minutes at 4 °C. Discard the pellet and centrifugation of the obtained supernatant at 100000 × g for 1 hour at 4 °C. The resulting pellet is ER fraction and the supernatant is cytosolic fraction. Resuspend pellet with lysis buffer (adding protease inhibitor cocktail) for subsequent WB analysis. For mouse brain mitochondria isolation, 6 months old wild type mouse was sacrificed after anesthetized and the brain was collected. The brain was removed and immediately rinsed three times with ice-cold start buffer (225 mM mannitol, 75 mM sucrose, 30 mM Tris–HCl, pH 7.4). The brain tissue was then minced and homogenized in isolation buffer (225 mM mannitol, 75 mM sucrose, 0.1 mM EGTA, 30 mM Tris–HCl, pH 7.4) at a ratio of 4 mL per gram of tissue using a Teflon-glass homogenizer with 20 up-and-down strokes. The homogenate was centrifuged at 700 × g for 5 minutes at 4°C. The supernatant was collected and centrifuged at 700 × g, 5 minutes, 4°C. The resulting supernatant was further centrifuged at 9000 × g for 10 minutes at 4°C. The pellet obtained represented the crude mitochondrial fraction and the supernatant was collected as previously described. After twice washing by start buffer, the crude mitochondrial was resuspended in MRB (250 mM mannitol, 5 mM HEPES, pH 7.4, 0.5 mM EGTA). The suspension of cell or mouse brain crude mitochondria was layered over Percoll medium (225 mM mannitol, 25 mM HEPES, pH 7.4, 1 mM EGTA, 30% Percoll) and centrifuged at 95,000 × g for 30 minutes at 4°C. The mitochondrial fraction (bottom layer) and MAM fraction (middle layer) were collected, resuspended in MRB at a 1:10 ratio, and centrifuged at 9000 × g for 10 minutes at 4°C twice. The mitochondria and MAM were lysed in cell lysis buffer (25 mM Tris-HCl, pH 7.6, 150 mM NaCl, 1% NP-40, protease inhibitor cocktail) for subsequent WB analysis. Protein concentrations of all fractions were determined by BCA assay for subsequent WB analysis. Wild-type C57BL/6JNifdc mice were obtained from Vital River (China). All experimental procedures were conducted in accordance with the animal ethics guidelines of Shandong University.

### Sub-Mitochondrial Fraction

For sub-mitochondrial compartment fractionation[32], purified mitochondria were resuspended in isolation buffer (225 mM mannitol, 75 mM sucrose, 0.1 mM EGTA, 30 mM Tris–HCl, pH 7.4) containing digitonin at a ratio of 0.12 mg digitonin per mg of mitochondrial and incubated with rotation for 15 min at 4□°C. Subsequently, a threefold volume of isolation buffer was added, and the mixture was centrifuged at 9000 ×□g for 10 min at 4□°C to yield an initial supernatant (Supernatant 1) and a pellet (Pellet 1). Supernatant 1 was ultracentrifuged under 144,000 ×□g, 1□h, 4 °C. The resulting pellet (Pellet 2) was lysed by appropriate volume of lysis buffer as outer mitochondrial membrane (OMM). The supernatant (Supernatant 2) was concentrated by TCA Protein Precipitation Kit (Sangon Biotech, China) as a inter membrane space protein (IMS). Pellet 1 was resuspended in isolation buffer and subjected to sonication on ice (3s/2s on/off, total 30s). The sonicated sample was centrifuged at 6500 ×□g for 10 min at 4□°C, and the resulting supernatant (Supernatant 3) was further ultracentrifuged at 144,000 ×□g for 1□h at 4□°C. The pellet (Pellet 4) representing the inner membrane fraction, was lysed in an appropriate volume of lysis buffer, while the supernatant (Supernatant 4) contained matrix proteins and was concentrated by TCA Protein Precipitation Kit (Sangon Biotech, China) for subsequent analysis.

### Mitochondrial Alkali assay

Crude mitochondria were treated with 0.1 M Na_2_CO_3_ on ice for 30 minutes. After centrifugation at 16,000 × g for 15 minutes at 4 °C, the supernatant (non-membrane fraction) was collected, and the pellet (mitochondrial membrane fraction) was washed with MRB and resuspended in cell lysis buffer for WB analysis.

### Mitochondrial Proteinase K Assay

Crude mitochondria were diluted to 1 μg/μL in MRB. Proteinase K (final concentration 800 ng/μL) was added with or without 0.2% Triton X-100 and incubated at room temperature for 15 minutes. The reaction was terminated by adding PSMF, and samples were analyzed by western blot.

### Western Blot (WB)

Cells were lysed in cell lysis buffer (25 mM Tris-HCl, pH 7.6, 150 mM NaCl, 1% NP-40, protease inhibitor cocktail) for 30 minutes at 4°C, followed by centrifugation at 16,000 × g for 10 minutes at 4°C. Protein concentration was determined by BCA assay. Samples were mixed with 5× loading buffer (250 mM Tris-HCl, pH 6.8, 10% SDS, 0.5% BPB, 50% glycerol, 5% β-mercaptoethanol), heated at 50°C for 8 minutes, and subjected to 10% SDS-PAGE. For ATAD3A oligomerization detection, samples were mixed with 5×loading buffer absence β-mercaptoethanol and analyzed by SDS-PAGE. Antibodies used in this study were listed in table 1.

**Table 1.** Antibodies used in this study.

| Antibody | Source | Identifier |
| --- | --- | --- |
| Rabbit anti-Flag | Thermo Fisher | PA1-984B |
| Rabbit anti-Myc tag | Bethyl Laboratories | A190-105A |
| Rabbit anti-Tim23 | Proteintech | 11123-1-AP |
| Rabbit anti-PS1-NTF | Cell Signaling Technology | 87146 |
| Rabbit anti-PS1-CTF | Abcam | ab76083 |
| Mouse anti-Calnexin | Abcam | ab22595 |
| Mouse anti-ACSL4 | Santacruz | sc-271800 |
| Mouse anti-Tubulin | Sigma | T5168 |
| Rabbit anti-ATAD3A | Abclonal | A8230 |
| Rabbit anti-MIC60 | Abclonal | A14107 |
| Rabbit anti-MFN2 | santacruz | sc-50331 |
| Rabbit anti-TFAM | Proteintech | 22586-1-AP |
| Mouse anti-OPA1 | BD bioscience | 612607 |
| Mouse anti-TOM20 | BD bioscience | 612278 |
| Mouse anti-Actin | Proteintech | 20536-1-AP |
| Mouse anti-DNA | Progene | 61014 |
| Rabbit anti-TOM70 | Proteintech | 14528-1-AP |
| Mouse anti-VDAC1 | Abcam | ab14734 |
| Mouse anti-IP3R-α | santacruz | sc-398434 |
| Mouse anti-cytochrome c | santacruz | sc-13156 |
| Peroxidase AffiniPure® Goat Anti-Mouse IgG (H+L) | Jackson ImmunoResearch Laboratories | 115-035-003 |
| Peroxidase AffiniPure® Goat Anti-Rabbit IgG (H+L) | Jackson ImmunoResearch Laboratories | 111-035-003 |
| CoraLite594-conjugated Goat Anti-Rabbit IgG(H+L) | Proteintech | SA00013-4 |
| CoraLite488-conjugated Goat Anti-Mouse IgG(H+L) | Proteintech | SA00013-1 |
| 12 nm Colloidal Gold AffiniPure® Goat Anti-Rabbit IgG (H+L) (EM Grade) | Jackson ImmunoResearch Laboratories | 111-205-144 |

### mtDNA Copy Number

Total cellular DNA was extracted using a genomic DNA extraction kit (Vazyme, China). Mitochondrial DNA was amplified using primers for the 12S gene (12s-F: 5′-TAGAGGAGCCTGTTCTGTAATCGA-3′, 12s-R: 5′-TGCGCTTACTTTGTAGCCTTCAT-3′), and nuclear DNA was amplified using primers for MLH1 (MLH1-F: 5′-GTAGTCTGTGATCTCCGTTT-3′, MLH1-R: 5′-ATGTATGAGGTCCTGTCCTA-3′). The mtDNA copy number was calculated using the 2^-ΔΔct^ method.

### DNA Repair Assay

Cells were treated with 150 μM H_2_O_2_ for 30 minutes at 37 □, and the mitochondrial DNA damage levels were quantified through comparative amplification efficiency analysis of long-range mtDNA fragments (10 kb), as described previously[33]. Total cellular DNA was extracted at various time points using a genomic DNA extraction kit (Vazyme, China). DNA damage and repair were assessed by qPCR using primers for large and small mitochondrial and nuclear fragments. The extent of DNA damage and repair was calculated by comparing the copy numbers of large fragments to small fragments using the 2^-ΔΔct^ method. qPCR was performed using SuperStar Universal SYBR Master Mix (Cwbio, China) on a Roche LC960 instrument. The primer nucleotide sequences used for the amplification of the 10 kb human mitochondrial DNA fragments are listed as follows: Large mitochondrial fragment Primer Sens: 5′-CCCACAGTTTATGTAGCTTACCTCCTCA-3′, Large mitochondrial fragment Primer Reverse: 5′-TTGATTGCTGTACTTGCTTGTAAGCATG-3′. The primers used for the amplification of the 172 bp human mitochondrial fragments are listed as follows: Small mitochondrial fragment Primer Sens: 5′-CTACACAATCAAAGACGCCC-3′, Small mitochondrial fragment Primer Reverse:5′-CTACACAATCAAAGACGCCC-3′, The primer used for the amplification of the nuclear DNA fragment are listed as follows: Large nuclear fragment Primer Sens : 5′-GAACGTCTTGCTCGAGATGTGATGAAGGAG-3′. Large nuclear fragment Primer Reverse: 5′-TCTCCCACCCATACTGGCAAAACTTAAGCC-3′. Small nuclear fragment Primer Sens: 5′-TCACATTGTAGCCCTCTGTG-3′, Small nuclear fragment Primer Reverse : 5′-ACACAATAGCTCTTCAGTCTG-3′.

### Immunoprecipitation (IP)

HEK293 cells, PS1KO cells, and PS1KO cells re-expressing PS1-wt were lysed in cell lysis buffer (25 mM Tris-HCl, pH 7.6, 150 mM NaCl, 1% NP-40, protease inhibitor cocktail) for 30 minutes at 4 °C, followed by centrifugation. The supernatant was incubated with the corresponding IP antibody overnight, then with Protein A/G magnetic beads (Selleck, China) for 2 hours at 4 °C. Beads were washed five times with cell lysis buffer, resuspended in 1× loading buffer, heated at 50 °C for 8 minutes, and analyzed by WB. To explore PS1-interacting proteins, Flag-tagged wild-type PS1 (Flag-PS1wt) immuneprecipitate and negative control were subjected to mass spectrometry (LC-MS/MS) for identification. The resulting data were then analyzed using Metascape to specifically profile mitochondrial proteins that interact with PS1.

### Transmission Electron Microscopy (TEM) and Immunoelectron Microscopy

For transmission electron microscopy, PS1 knock out cells re-expressing Flag tagged wild type PS1 were collected by centrifugation, fixed in electron microscopy fixative (Servicebio G1102, China) at 4 °C, washed with 0.1 M phosphate buffer (pH 7.4), embedded in 1% agarose, and post-fixed with 1% OsO_4_ in 0.1 M phosphate buffer (pH 7.4) for 2 hours. washed, and dehydrated through a graded ethanol series (30%, 50%, 70%, 80%, 95%, and 100%) followed by two changes of acetone. Samples were infiltrated with EMBed□812 resin using graded acetone□resin mixtures (1:1 and 1:2) and pure resin, embedded in pure resin, and polymerized at 60□°C for 48□h. Ultrathin sections (60–80□nm) were collected on Formvar□coated 150□mesh copper grids, stained with 2% uranyl acetate and 2.6% lead citrate. Samples were imaged using a HITACHI HT7800/HT7700 transmission electron microscope.

For immuno-electron microscopy, PS1 knock out cells re-expressing Flag tagged wild type PS1 were collected, fixed in immunoelectron microscopy fixative (Servicebio G1124, China) at 4□°C in the dark, and washed with ice□cold 0.1□M phosphate buffer (PB, pH□7.4). The fixed cells were embedded in 2% low□melting□point agarose (Solarbio, China), dehydrated through a graded ethanol series (30% to 100%) at 4□°C to □20□°C, and infiltrated with LR□White resin (HaideBio, China) using graded mixtures of ethanol and resin, followed by pure resin changes. Samples were embedded in capsules, degassed under vacuum, and polymerized under UV light at □20□°C for >48□hours. Ultrathin sections (70□80□nm) were collected on formvar□coated nickel grids, blocked with 1% BSA, and incubated overnight at 4□°C with primary anti-Flag (Thermo, PA1984B, USA) antibody diluted (1:100) in 1% BSA/TBS, followed by gold□conjugated secondary antibody (Jackson 111-205-144, USA, 1:25 diluted) incubation (20□min at RT, then 1□hour at 37□°C). After washing, grids were stained with 2% uranyl acetate (8□minutes, dark), dried, and examined by TEM, with 12□nm gold particles indicating specific labeling.

### Seahorse Assay

Mitochondrial oxygen respiration capacity was assessed using the XF Cell Mito Stress Test Kit (Agilent, USA). 1500 cells were seeded in XFp 24-well microplates with 250 μL of growth medium (DMEM + 10% FBS + 1% P-S) per well and cultured overnight. Oxygen consumption rate (OCR) was measured using the Agilent Seahorse XFe24 instrument, with sequential additions of oligomycin (1.5 μM), FCCP (1.5 μM), and rotenone/antimycin A (0.5 μM). After test cells in each well were lysed on ice for 15 minutes using lysis buffer. The protein concentration of each lysate was determined by BCA assay, and the obtained oxygen consumption rate (OCR) values were normalized to the total protein content.

### Peptide inhibitor

To inhibit ATAD3A oligomerization, we synthesized a peptides inhibitor DA1(GRKKRRQRRRPQ-GG-EDKRKT-NH_2_) and comparative control peptides TAT(GRKKRRQRRRPQ-GG-NH_2_) [29]. Peptides were synthesized at Genechem (China). The purity was assessed as >90% by mass spectrometry. PS1KO cells cultured with 1μM DA1 at least 48h to decrease ATAD3A oligomerization.

### Statistical analysis

All results were presented as mean ± standard deviation. Statistical analyses were performed using GraphPad Prism 8.0 software. Student’s t-test was used for comparisons between two groups, whereas one-way analysis of variance (one-way ANOVA) was applied for comparisons among multiple groups. A p-value < 0.05 was considered statistically significant.

## RESULTS

### Presenilin 1 (PS1) Localizes to the Mitochondrial Inner Membrane

PS1, a nine-transmembrane protein with self-cleavage activity, is primarily localized in the endoplasmic reticulum (ER) and mitochondria-associated membrane (MAM). Although earlier studies have reported the presence of PS1 on the mitochondrial membrane, its specific localization-whether on outer or inner mitochondrial membrane-remains unclear. To address this, we isolated mitochondria, MAM, and ER membrane fractions from mouse brain and PS1 knock out (KO) cells re-expressing Flag-PS1wt using density gradient centrifugation, and assessed PS1 distribution across these fractions. As shown in Figure 1A and 1B, PS1 was primarily detected in ER and MAM fractions. Moreover, the full length PS1, N-terminal PS1 and C-terminal PS1 could also be detected in mitochondrial fractions (Fig 1A, 1B). Confocal microscopy further confirmed partial co-localization of PS1 with TIM23 (Translocase of Inner Mitochondrial Membrane 23), a marker of the mitochondrial inner membrane (Fig. 1C). To further delineate the submitochondrial localization of PS1, we performed an alkali assay, which collected mitochondrial membrane and non-membrane fractions, confirming PS1 distribution in the mitochondrial membrane fraction (Fig. 1D). Additionally, proteinase K digestion assays on cell mitochondria revealed that PS1 could not be digested by proteinase K when the mitochondrial membrane structure is intact, but be digested in the presence of both Triton-X 100 and proteinase K (Fig. 1E), supporting its localization on the mitochondrial inner membrane. To definitively establish the submitochondrial localization of PS1, we performed organelle fractionation, which verified its specific residency within the inner mitochondrial membrane (Fig. 1F). Finally, through immunoelectron microscopy, we directly visualized PS1 on the mitochondrial inner membrane (Fig. 1G-I). Collectively, these above results suggested that PS1 could localize on mitochondrial inner membrane.

**Fig. 1.**
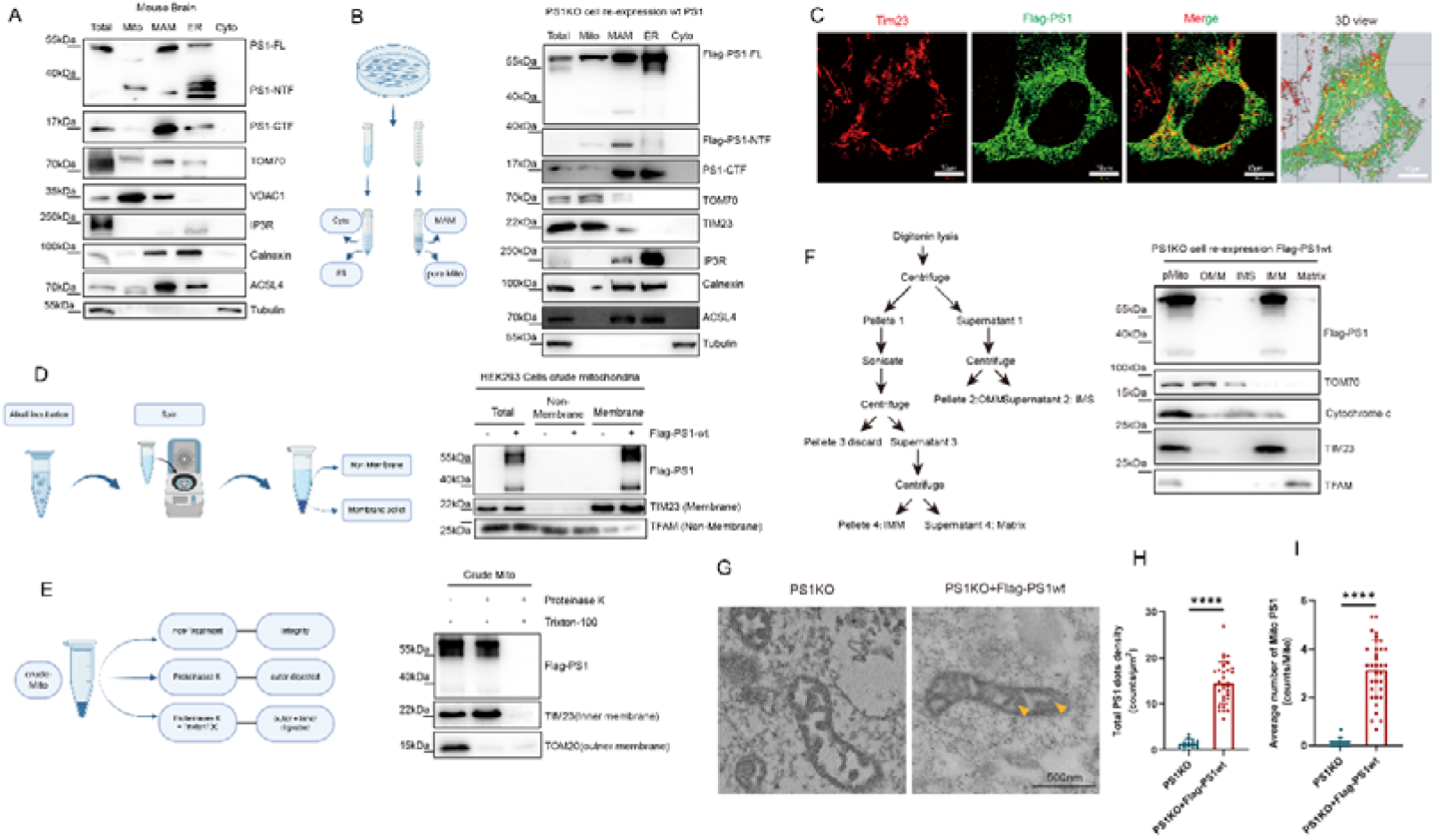
PS1 located at mitochondrial inner membrane. **A and B** Isolated sub-cellular compartments from mouse brain**(A)** or HEK293 cells**(B)** were analyzed by WB to detect PS1 location. Total: total cell lysate; Mito: pure mitochondrial; MAM: mitochondrial associated membrane; ER: endoplasmic reticulum; Cyto: Cytosol. Markers: ACSL4 (MAM), IP3R & Calnexin (ER), VDAC1, TOM70 & TIM23 (Mitochondria), Tubulin (Cytosol). **C** Immunofluorescence shows colocalization of PS1 and mitochondrial inner membrane protein TIM23. Scale bar: 20μm. **D** Crude mitochondria were prepared and subjected alkali extraction and western blot to analyze mitochondrial membrane and non-membrane proteins. **E** Crude HEK293 cells mitochondrial were incubated with proteinase *K* in presence or absence Triton X-100 before immunoblot. **F** WB analysis of sub-mitochondrial fractions indicates the inner membrane localized PS1. Left: Schematic diagram of submitochondrial fractionation by digitonin lysis. Right: pMito, pure mitochondria; OMM, outer mitochondrial membrane; IMS, intermembrane space; IMM, inner mitochondrial membrane. Markers: TOM70 (OMM), Cytochrome□c (IMS), TIM23 (IMM), and TFAM (matrix). **G** Immuno-electron microscope shows PS1 located at mitochondrial inner membrane. The arrow indicates the PS1 golden particles. Images were acquired at 25,000× magnification. Scale bar: 500nm. **H** Total density of PS1□specific immunogold particles (counts per μm²). **I** Average number of PS1□labeled gold particles per mitochondrion (mitochondrial PS1 dots per mitochondrion). A total of 16 fields of view from PS1KO cells and 30 fields of view from PS1KO + Flag□PS1wt cells were analyzed. Data was presented as the mean ± SD. ⍰*p* <0.05. ⍰⍰*p* <0.01. ⍰⍰⍰*p* <0.005. ⍰⍰⍰⍰*p* <0.001. n.s. no significant.

### PS1 deficiency impairs mitochondrial function

Having established the submitochondrial localization of PS1, we next sought to determine its functional relevance. Mitochondria are essential organelles in eukaryotic cells, playing central roles in energy metabolism, reactive oxygen species (ROS) production, and calcium (Ca^2+^) homeostasis regulation[15, 16, 18, 19, 34]. Having demonstrated the localization of PS1 on the mitochondrial inner membrane, we next evaluated the impact of PS1 on mitochondrial function. PS1 knockout HEK293 cell lines were generated using CRISPR-Cas9 and confirmed PS1 protein absence by WB (Fig. 2A-B). We evaluated cell proliferation using CCK8 assays and found significant reduced proliferation of PS1KO cells (Fig. 2C). Using the Seahorse, we observed that PS1 deficiency significantly reduced the basal oxygen respiration rate (OCR) and ATP synthesis, while increasing proton leakage (Fig. 2D-G). PS1KO cells displayed significantly decrease in mitochondrial membrane potential, which was rescued by re-expressing wildtype (wt) PS1 (Fig. 2H and 2I). Using mitochondrial-targeted Ca^2+^ and ROS probes, we observed elevated mitochondrial Ca^2+^ levels and enhanced mitochondrial ROS generation (Fig. 2J and 2K, 2L) in PS1KO cells, indicating disrupted Ca^2+^ homeostasis and ROS production. While re-expression of wt PS1 in PS1KO cells attenuated these abnormalities in Ca^2+^ and ROS. In summary, PS1 deficiency compromises mitochondrial bioenergetic metabolism, resulting in suppressed cell proliferation, reduced ATP production, disrupted mitochondrial calcium homeostasis, and excessive ROS accumulation. These findings underscore the role of PS1 in regulating mitochondrial physiology.

**Fig. 2.**
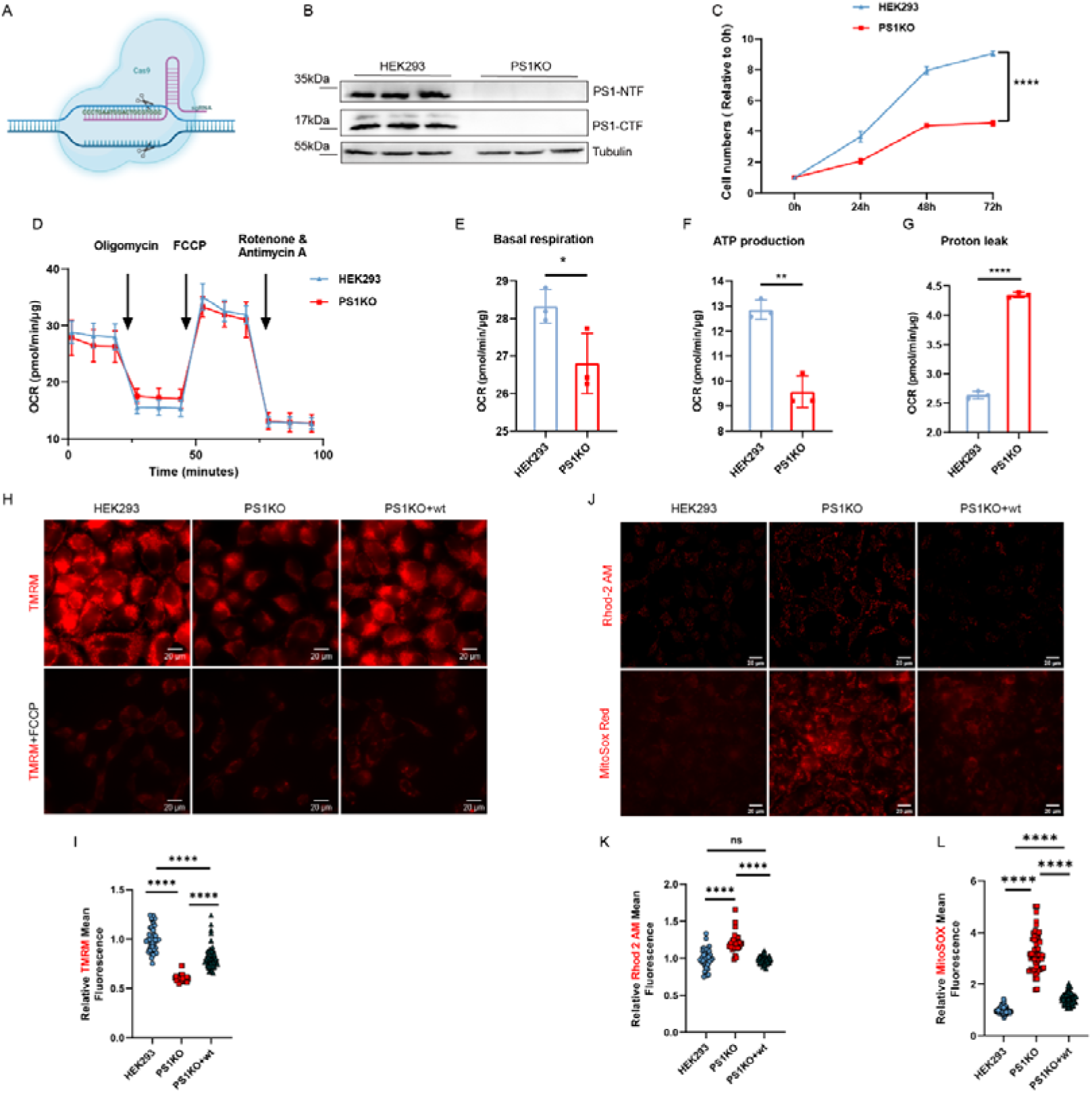
PS1 deficiency impair mitochondrial function. **A** Schematic of CRISPR/Cas9 genome-editing of the PS1 gene in HEK293 cells to obtain PS1KO cells. **B** PS1 knock out cells validated by western blot. **C** PS1KO cells proliferation capacity was detected by Cell Counting Kit-8. **D** Mitochondrial Oxidative respiration capacity was analyzed by Seahorse XF cell mitochondrial stress test. Quantification of OCR normalized to total protein of each well. The Basal respiration(**E**) and ATP production(**F**) was decreased in PSKO cell, but the Proton leak(**G**) was increased in PS1KO cell. **H, I** Mitochondrial potential was analyzed by TMRM staining and living cell imaging. As a negative control cell were incubated with FCCP before TMRM staining. The mean fluorescence of TMRM was quantified and normalized to HEK293 group. HEK293 *n* = 346 cells (43 fields of view). PS1KO *n* = 368 cells (41 fields of view). PS1KO+wt *n* = 446 cells (56 fields of view). Scale bar: 20μm. **J, K** Mitochondrial Ca^2+^ level was analyzed by Rhod-2 AM staining and living cell imaging. The mean fluorescence of Rhod-2 AM was quantified and normalize to HEK293 group. HEK293 *n* = 336 cells (37 fields of view). PS1KO *n* = 307 cells (34 fields of view). PS1KO+wt n = 351 cells (43fields of view). Scale bar: 20μm. Scale bar: 20μm. **J, L** Reactive Oxygen Species (ROS) level was analyzed by MitoSOX staining and living cell imaging. The fluorescence of MitoSOX was quantified and normalize to HEK293 group. HEK293 *n* = 713 cells (73 fields of view). PS1KO *n* = 563 cells (54 fields of view). PS1KO+wt *n* = 619 cells (67 fields of view). Scale bar: 20μm. Data was presented as the mean±SD. ⍰*p* <0.05. ⍰⍰*p* <0.01. ⍰⍰⍰*p* <0.005. ⍰⍰⍰⍰*p* <0.001. n.s. no significant.

### Loss of PS1 leads to abnormal mitochondrial morphology and vacuolation

Maintenance of functional competence in mitochondria depends on their structure integrity [19]. To determine the effects of PS1 deficiency on mitochondrial structural consequences, we investigated the mitochondrial morphology in PS1 KO cells using Super-Resolution imaging with inner membrane-targeted fluorescent probes. The results demonstrated that mitochondria in PS1 KO cells exhibited less branched networks and vacuolar structures (Fig. 3A-3D). Quantitative analysis revealed a significant reduction in total mitochondrial area per cell (Fig. 3B), and mitochondrial aspect ratio (Fig. 3E). And re-expression of wt PS1 could restore these morphological defects (Fig 3 A-E). We then performed ultrastructural and morphological analyses using transmission electron microscopy, which revealed pronounced abnormalities in mitochondrial inner membrane of PS1 KO cells, including a decreased number of cristae and swollen cristae morphology (Fig. 3F-I). Together, these findings demonstrate that PS1 deficiency disrupts both global mitochondrial architecture and ultrastructure of the inner membrane, thereby providing structure basis for the accompanying functional impairments.

**Fig. 3.**
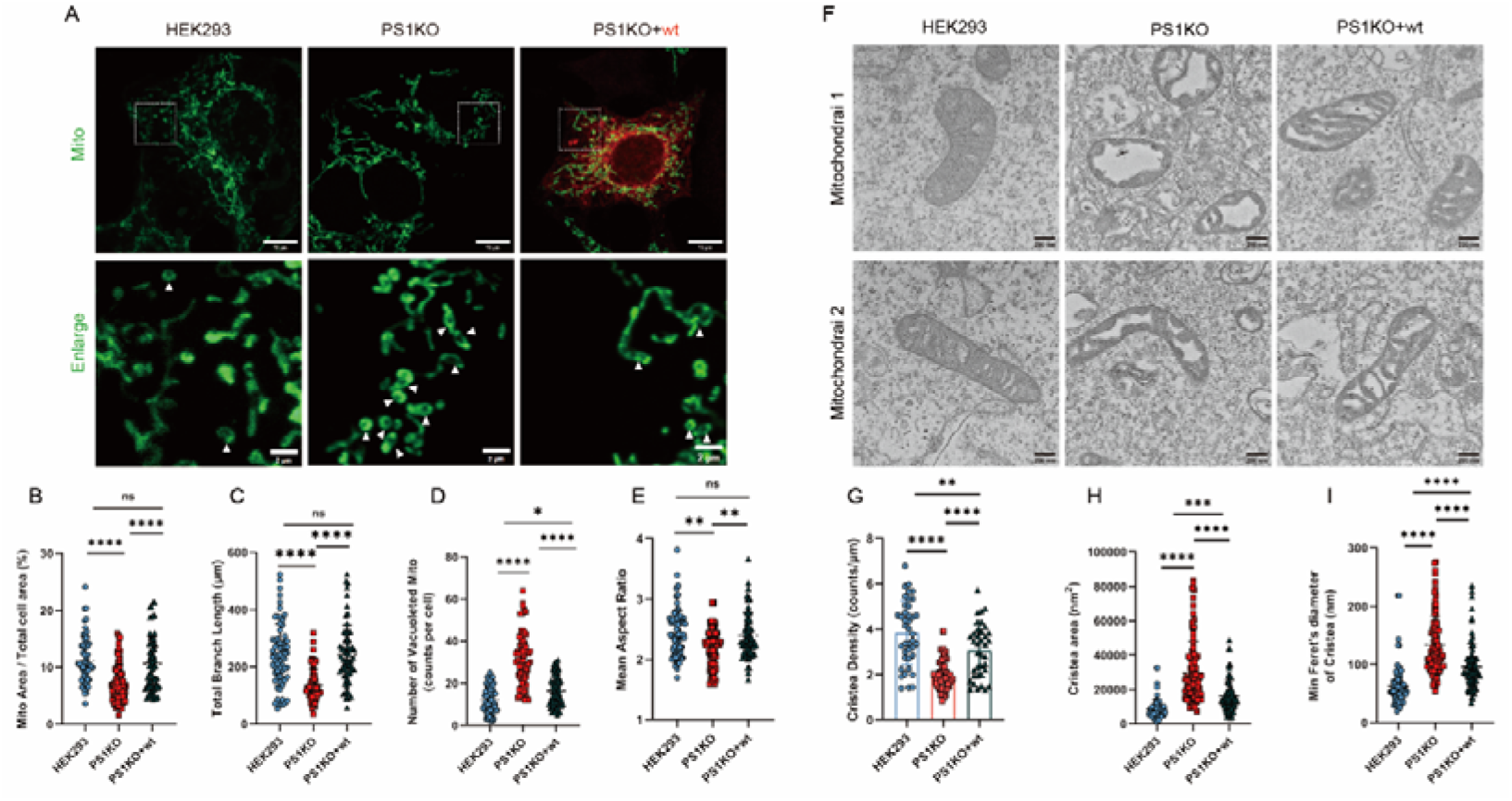
Loss of PS1 induced mitochondrial vacuolation and fragment. **A** Mitochondrial morphology was analyzed by Mito-EGFP labeling and Zeiss LSM900 airyscan2. White arrows indicate the vacuolated mitochondrial. **B-D** Quantification of mitochondrial morphology. **B** Percentage of mitochondrial area relative to total cell area per cell. **C** total branch length of single cell mitochondria. **D** Number of vacuolated mitochondria per cell. **E** mean aspect ratio of cell mitochondria. *n*= 58 HEK293 cells, *n*= 76 PS1KO cells, *n*= 70 PS1KO + wt cells were analyzed. **F** Transmission electro-microscope indicated that PS1KO cell mitochondrial inner membrane disruption. The upper and lower panels are from the same experiment; the lower image shows that, although tubular mitochondria are still present in PS1□KO cells, they exhibit damaged cristae architecture. **G** Cristae density of mitochondria (number of cristae per mitochondrial perimeter). **H** Mitochondrial cristae area is increased in PS1KO cells. **I** The Min Feret’s diameter of cristae is elongated in PS1□KO mitochondria, indicating a swollen cristae morphology. *n*= 40 HEK293 cell mitochondria, *n*= 43 PS1KO cell mitochondria, *n*= 37 PS1KO + wt mitochondria were analyzed. Data was presented as the mean ± SD. ⍰*p* <0.05. ⍰⍰*p* <0.01. ⍰⍰⍰*p* <0.005. ⍰⍰⍰⍰*p* <0.001. n.s. no significant.

### Loss of PS1 induced mitochondrial inner membrane protein ATAD3A oligomerization

To explore the molecular mechanism underlying mitochondrial inner membrane damage upon PS1 deficiency, we identified PS1-interacting proteins through immunoprecipitation coupled with mass spectrometry to. Gene Ontology (GO) analysis revealed a number of interactors are mitochondria-localized proteins (Fig. 4A), with the majority residing in the inner membrane (Fig. 4B-D). Since mitochondrial membrane proteins are essential for cristae arrangement, we selected several for further validation. Co-immunoprecipitation (co-IP) confirmed that PS1 interacts with the mitochondrial inner membrane protein OPA1, MIC60, ATAD3A and TIM23, as well as the mitochondrial matrix protein TFAM (Fig. 4C), all of which contribute to the maintenance of mitochondrial inner membrane architecture. In contrast, PS1 did not interact with the outer mitochondrial membrane proteins TOM70 and MFN2 (Fig. 4C). We next assessed whether PS1 deficiency alters mitochondrial protein expression. Western blot analysis revealed that loss of PS1 led to decreased MFN2, and TIM23, but increased ATAD3A in total cell lysates. (Fig. 4D-E). ATAD3A is a mitochondrial membrane protein with its C-terminus in the mitochondrial inner membrane and N-terminus spanning the outer membrane. It interacts with MICOS complex and TFAM to maintain normal inner membrane structure and mtDNA stability[22, 24, 28]. In AD models, ATAD3A level changes impair mitochondrial and MAM function[28, 29]. We further focused on ATAD3A and found PS1 deficiency not only increased its protein level but also promoted its oligomerization, as detected by SDS-PAGE under non-reducing conditions (without β-mercaptoethanol) (Fig. 4F-H). ATAD3A oligomerization has been implicated mitochondrial dysfunction in AD, suggesting mitochondrial damage induced by PS1 deficiency may be associated with increased ATAD3A level and oligomerization.

**Fig. 4.**
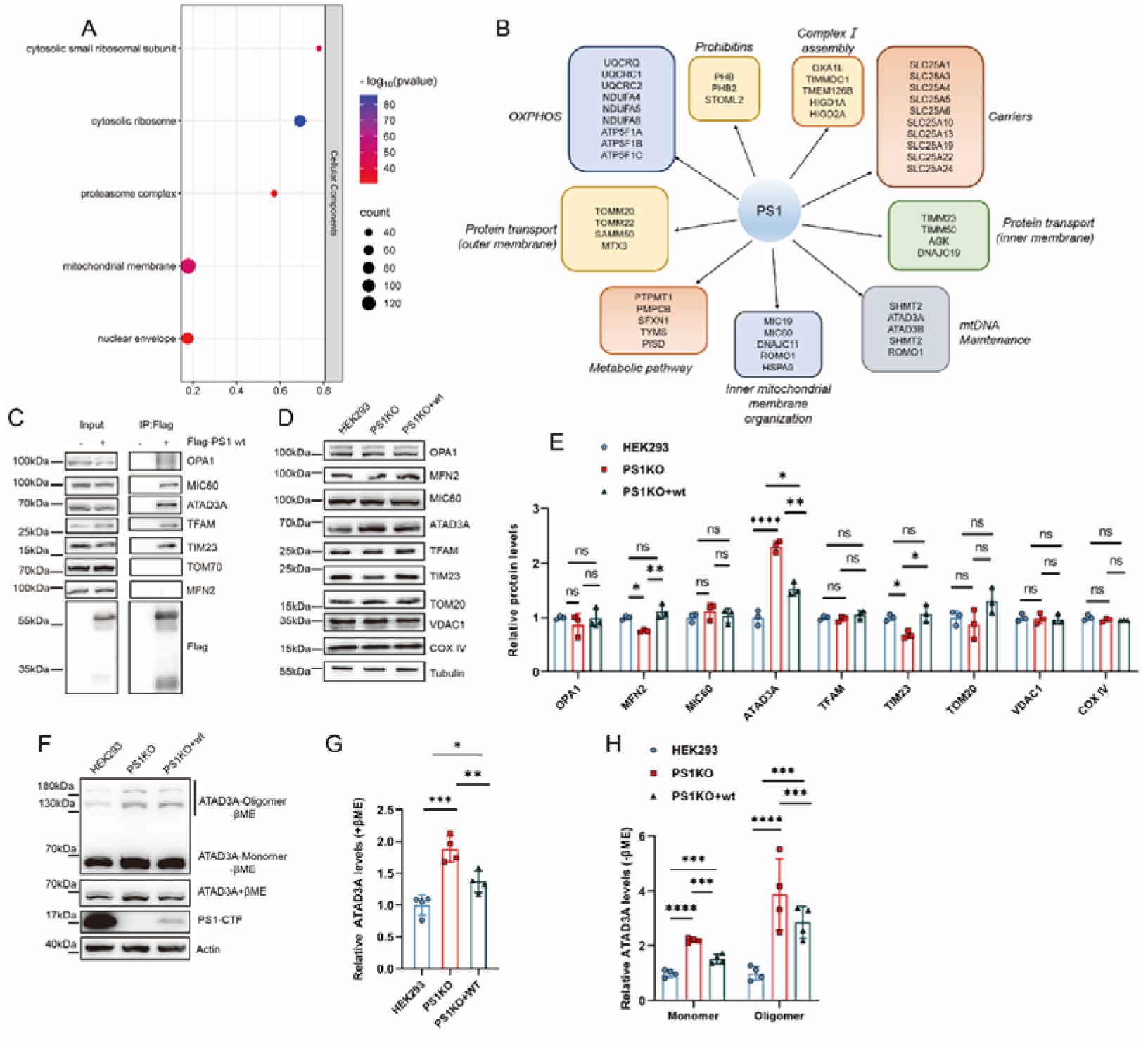
Loss of PS1 induced mitochondrial inner membrane protein ATAD3A oligomerization. **A-B** GO-cell compartment analysis indicated that mitochondrial membrane protein interacted with PS1. **C** co-immunoprecipitation confirmed the interaction between PS1 and mitochondrial inner membrane and matrix protein, but not with outer mitochondrial membrane protein TOM70 or MFN2. **D** Analysis of mitochondrial protein alterations in total cell lysates by Western blot. **E** Quantification of mitochondrial protein levels in total cell lysate. **F** ATAD3A protein levels in total cell lysate were determined by western blotting (WB) with anti-ATAD3A antibody in the presence or absence of β-mercaptoethanol (βME). **G-H** Quantification of ATAD3A monomer and oligomer level. Data was presented as the mean ± SD. ⍰*p* <0.05. ⍰⍰*p* <0.01. ⍰⍰⍰*p* <0.005. ⍰⍰⍰⍰*p* <0.001. n.s. not significant.

### ATAD3A oligomerization in PS1 knock out cells leads to expanded MAM area and mtDNA repair dysfunction

Given that ATAD3A oligomerization have been shown to promote the expansion of MAM structures, and disrupt mitochondrial and ER function[22, 27, 35], we asked whether MAM structure is altered in PS1 KO cells. Using the SPLICS Mt-ER Long P2A plasmid for labeling, we observed a significant increase in MAM area in PS1 KO cells (Fig. 5A-B). Since mtDNA integrity is closely linked to mitochondrial structure, we also examined mtDNA copy number and found it to be significantly reduced in PS1 KO cells mtDNA copy numb (Fig. 5C). Consistent with this, immunostaining with anti-DNA antibody revealed fewer mtDNA nucleoids in PS1KO cells (Fig. 5D-E).

**Fig. 5.**
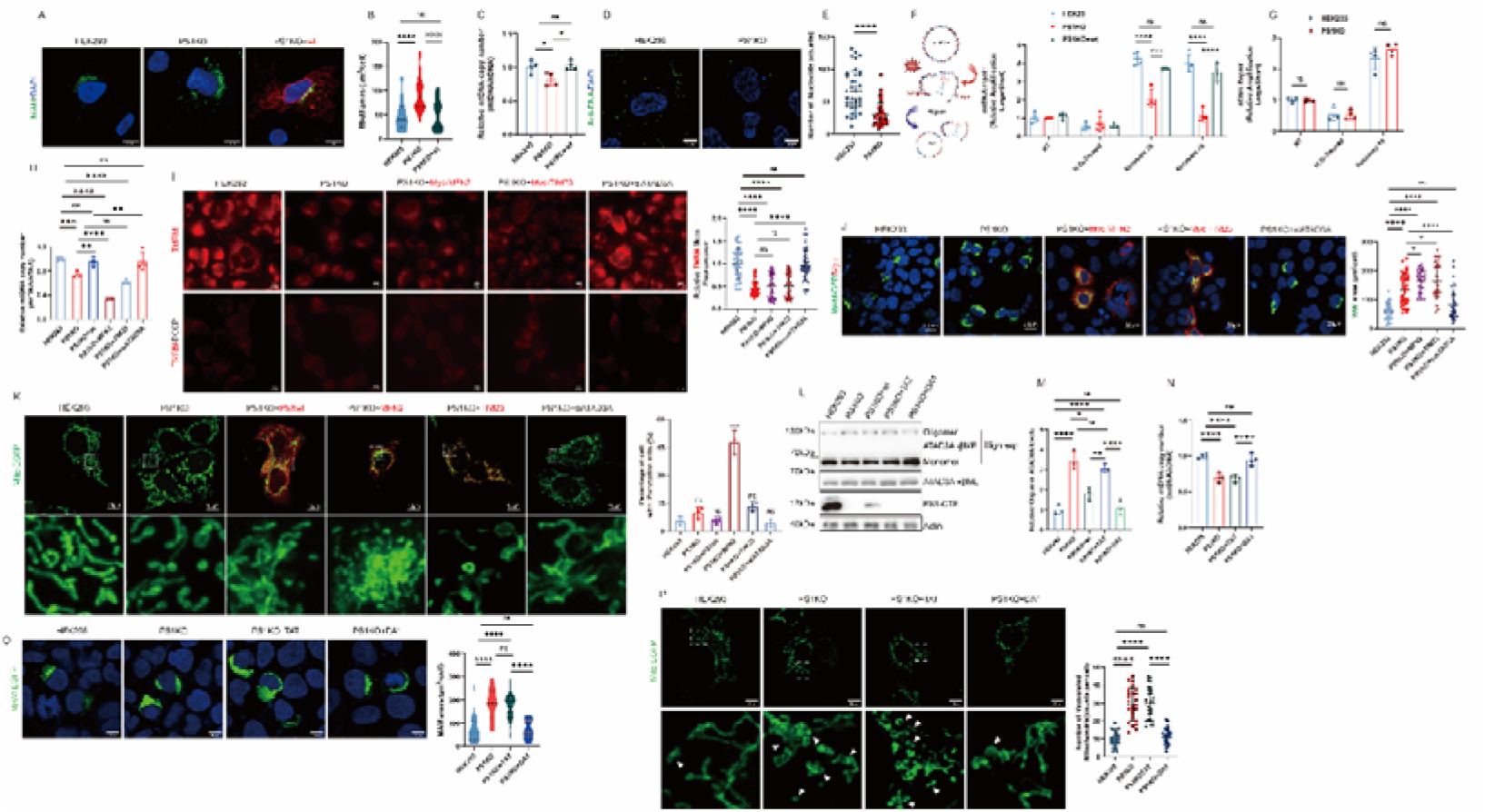
ATAD3A oligomerization in PS1 knock out cells lead to expanded MAM area and mtDNA repair dysfunction. **A** Mitochondrial associated membrane (MAM) area was determined by SPLICS Mt-ER *L* plasmid. **B** Quantification of MAM area. n= 32 cells were analyzed. **C** Mitochondrial DNA copy number was analyzed by quantitative PCR. **D** Mitochondrial DNA nucleoids were determined by anti-DNA staining. **E** Anti-DNA puncta were quantified. HEK293 n=37, PS1KO n=40 cells were analyzed. **F** Mitochondrial DNA repair assay by using quantitative PCR. Left: Schematic of H_2_O_2_ induced DNA disruption and repair. **G** Nuclear DNA repair assay by using quantitative PCR. **H** Quantification of mtDNA copy number in PS1KO cells following overexpression of MFN2 or TIM23, or knockdown of ATAD3A. **I** Measurement of mitochondrial membrane potential by TMRM staining in PS1KO cells following overexpression of MFN2 or TIM23, or knockdown of ATAD3A. Right: Quantification of relative TMRM fluorescent. TMRM fluorescent were normalized to HEK293. HEK293 *n*=720 cells (40 fields of view), PS1KO *n*=644 cells (56 fields of view), PS1KO+MFN2 *n*=506 cells (46 fields of view), PS1KO+TIM23 *n*=564 cells (47 fields of view), PS1KO+siATAD3A *n*=675 cells (45 fields of view). **J** MAM area was determined by SPLICS Mt-ER *L* plasmid and quantified in PS1KO cells following overexpression of MFN2 or TIM23, or knockdown of ATAD3A. Right: Quantification of MAM area. *n*= 40 HEK293 cells, *n*=44 PS1KO cells, *n*=40 PS1KO+MFN2 cells, *n*=41 PS1KO+TIM23 cells *n*=42 PS1KO+siATAD3A cells were analyzed. **K** Mitochondria in PS1KO cells following overexpression of MFN2 or TIM23, or knockdown of ATAD3A labeling by Mito-EGFP plasmids. Right: Quantification of percentage of cell with punctation Mitochondria. Data are representative of three independent experiments. **L** PS1KO cells were treated with DA1 peptides or control peptides TAT (1μM each) for 48h. ATAD3A oligomerization were analyzed by WB. **M** Quantification of ATAD3A oligomerization levels. **N** Mitochondrial DNA copy number of peptides treated PS1KO cells was analyzed by quantitative PCR. **O** PS1KO cells were treated with peptides (DA1/TAT, 1μM each) for 48h, MAM area were labeling by transfected with SPLICS Mt-ER L plasmid. Right: Quantification of MAM area. *n*= 25 cells were analyzed. **P** PS1KO cells were treated with peptides (DA1/TAT, 1μM each) for 48h, the mitochondrial were labeling by transfected with Mito-EGFP plasmids. Right: Quantification of vacuolated mitochondrial. *n*= 25 cells were analyzed Data was presented as the mean ± SD. ⍰*p* <0.05. ⍰⍰*p* <0.01. ⍰⍰⍰*p* <0.005. ⍰⍰⍰⍰*p* <0.001. n.s. not significant.

We next evaluated the sensitivity of PS1 KO cells to oxidative stress. Following H_2_O_2_-induced DNA damage, PS1KO cells exhibited defective mtDNA repair (Fig. 5F), whereas nuclear DNA repair was unaffected and completed within one hour of recovery (Fig. 5G). Re-expression of wild type PS in PS1 KO cells restored MAM and mtDNA integrity to levels comparable to those in wildtype HEK293 cells (Fig. 5A-G).

To determine whether the mitochondrial structural defects, MAM expansion, and mtDNA damage in PS1KO cells are caused by the decrease in MFN2/TIM23 or by the elevation and oligomerization of ATAD3A, we performed rescue experiments by overexpressing MFN2 or TIM23 and by knocking down ATAD3A with siRNA in PS1KO cells. We first found that overexpression of MFN2 or TIM23 in PS1KO cells induced severe cytotoxicity and reduced cell viability, whereas ATAD3A knockdown had no obvious effect on survival (Fig. S1A-F). Correspondingly, it did not rescue the decreased mtDNA copy number or restore the loss of mitochondrial membrane potential (Fig. 5H-I). In contrast, ATAD3A knockdown in PS1KO cells significantly increased mtDNA copy number and recovered mitochondrial membrane potential (Fig. 5H-I). Furthermore, ATAD3A knockdown markedly reduced the expanded MAM area, while overexpression of MFN2 or TIM23 even further enlarged it (Fig. 5J). Together, these results demonstrate that the mitochondrial structural damage, MAM expansion, and mtDNA impairment in PS1KO cells are primarily driven by upregulation and oligomerization of ATAD3A, rather than by the reduction of MFN2 or TIM23.

Considering the effects of ATAD3A oligomerization on MAM regulation and mtDNA maintenance[22, 24, 28, 29, 36], we hypothesized that PS1 deficiency impairs mitochondrial functions through increased level and oligomerization ATAD3A. To confirm this, we generated a peptides DA1, corresponding to the residues 113-118aa of C-terminal region of ATAD3A, which interacts with ATAD3A colied-coli domain and decrease its oligomerization [28, 29]. Treatment with DA1, but not the control TAT peptide, significantly decreased oligomerization of ATAD3A in PS1KO cells (Fig. 5L-M), and also restored mtDNA copy numbers (Fig. 5N). Furthermore, DA1 treatment also restore the mitochondrial morphology and normalized MAM area in PS1KO cells (Fig. 5O and 5P).

Together, these findings indicate that PS1 deficiency could increase oligomerization of ATAD3A, which in turn drives the MAM expansion area and directly compromises mitochondrial structure and mtDNA maintenance.

### Loss of PS1 disturb mitochondrial cristae junction proteins arrangement

The maintenance of intact cristae structure requires proper assembly of protein complex in the mitochondrial inner membrane, such as MICOS complex, and the interaction between MICOS and other proteins such as OPA1 and ATAD3A [19, 22, 37]. To investigate the mechanism by which ATAD3A oligomerization impairs mitochondrial function in PS1KO cells, we further performed co-IP to assess the interactions between ATAD3A and component of the MICOS complex, OPA1 and other partners in both PS1KO and PS1-re-expressed cells. We found that PS1 deficiency increased the interactions of ATAD3A with MIC60, MIC19 and OPA1 (Fig. 6A-F), while leaving the ATAD3A-ATPase ATP5A interaction unchanged (Fig. 6G-H). Re-expression of wt PS1 restored these interactions to normal level (Fig. 6A-F). Similarly, DA1 treatment, which ameliorates mitochondrial and MAM dysfunction in PS1KO cells, also normalized the exaggerated interaction between ATAD3A and Mic60, Mic19, and OPA1 (Fig. 6I-J). Based on these results, we propose a model in which mitochondrial inner membrane localized-PS1 coordinates proper protein arrangement. Loss of PS1 leads to increased ATAD3A levels and oligomerization, which in turn disrupts its physiological binding with cristae junction proteins such as Mic60, Mic19, and OPA1, thereby compromising the structural integrity of the mitochondrial inner membrane (Fig. 6K).

**Fig. 6.**
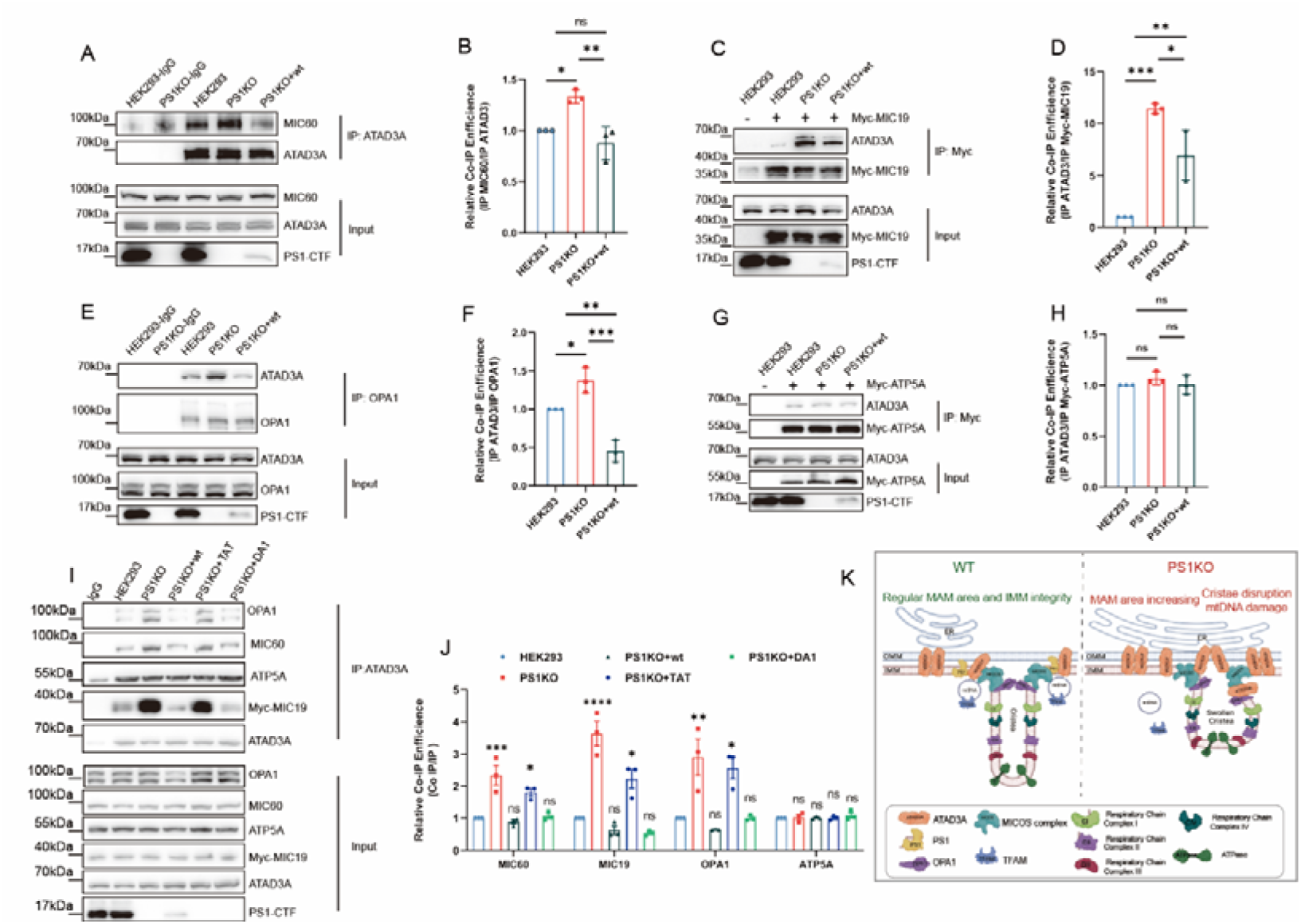
Loss of PS1 disturb mitochondrial cristae junction proteins arrangement. **A-F** The interaction between ATAD3A and mitochondrial cristae proteins were determined by co-immunoprecipitation. **A** The increased interaction between ATAD3A and MICOS proteins MIC60 in PS1KO cell. **B** Quantification of binding efficiency between ATAD3A and MIC60. **C** The increased interaction between ATAD3A and MICOS proteins MIC19 in PS1KO cell. **D** Quantification of binding efficiency between ATAD3A and MIC19. **E** The increased interaction between ATAD3A and OPA1 in PS1KO cell. **F** Quantification of binding efficiency between ATAD3A and MIC60. **G** The interaction between ATAD3A and ATPase was not changed. **H** Quantification of binding efficiency between ATAD3A and ATAD3A. **I** PS1KO cells were treated with DA1/TAT peptides for 72h(DA1/TAT, 1μM each), the interaction between ATAD3A and MIC60, OPA1, MIC19, ATP5A was analyzed by co-immunoprecipitation. **J** Quantification of binding efficiency between ATAD3A and mitochondrial membrane protein. **K** A summary scheme shows that loss of PS1 disturb mitochondrial cristae junction proteins arrangement and caused swollen cristae. Data are presented from three independent biological replicates of co-immunoprecipitation. For each replicate, the co-IP efficiency was calculated relative to the control HEK293 cells. Data was presented as the mean ± SD. ⍰*p* <0.05. ⍰⍰*p* <0.01. ⍰⍰⍰*p* <0.005. n.s. not significant.

## Discussion

In this study, we demonstrated that PS1 localizes to the mitochondrial inner membrane and regulates the arrangement of cristae junction architecture. Loss of PS1 induced mitochondrial vacuolization and dysfunction, characterized by reduced mitochondrial membrane potential, impaired oxidative phosphorylation, and decreased ATP production, accompanied by mitochondrial calcium overload and ROS accumulation. Further mechanistic investigation revealed that these structural and functional deficits arise from upregulation and oligomerization of the key mitochondrial protein ATAD3A. Specifically, PS1 knockout elevated ATAD3A protein expression, promoted its oligomerization, expanded MAM areas, and compromised mtDNA stability. These findings uncover a γ-secretase-independent role of PS1 in modulating mitochondrial morphology and function, potentially representing a critical mechanism underlying mitochondrial dysfunction in AD pathogenesis.

A key finding of this study is that PS1 localizes to the mitochondrial inner membrane and plays an important role in maintaining cristae morphology. Although mitochondrial localization of PS1 has been previously suggested[11], we provide substantial evidence that a fraction of PS1 resides specifically in the mitochondria inner membrane. We propose that PS1, ATAD3A and MICOS form a supercomplex within the inner membrane. In this model, inner membrane localized PS1 may help stabilize ATAD3A, thereby contributing to cristae junction integrity. Loss of PS1 could release ATAD3A from the inner membrane, promoting its oligomerization at the outer membrane and MAM regions, and enhancing its interact with MIC60 and Mic19-core components of the MICOS complex (Fig. 6). ATAD3A, which spans both mitochondrial membranes, is a known regulator of mitochondrial inner membrane dynamics[24, 26, 27]. Reduced inner membrane retention of ATAD3A and its aberrant binding to MICOS components likely disrupt cristae junction architecture and inner membrane morphology.

It has been demonstrated that ATAD3A participates in diverse mitochondrial functional processes, including nucleoid organization, cholesterol metabolism, and mitochondrial translation[25] For instance, it interacts with mitochondrial transcription factor A (TFAM) to regulate nucleoid dynamics and mtDNA transcription, while also mediating respiratory chain complex assembly[22, 36]. ATAD3A is also functionally linked to mitochondria-associated ER membranes (MAM), and regulates mitochondrial Ca^2+^ homeostasis [24, 29, 38]. Thus, the disrupted mitochondrial Ca^2+^ handling and mtDNA maintenance observed in PS1-deficiency cells stem from increased ATAD3A levels and oligomerization. These results underscore the central role of ATAD3A in mediating mitochondrial dysfunctions induced by PS1 deficiency. Notably, elevated ATAD3A oligomerization has been reported in AD patients and models, highlighting its potential pathogenic relevance [28, 29]. The precise mechanisms by which PS1 regulates ATAD3A, however, requires further investigation.

Our study also reveals a novel γ-secretase-independent functions of PS1, which may contribute to AD progression. PS1 undergoes autoproteolytic processing to form N-terminal (NTF) and C-terminal (CTF) fragments, which together constitute the catalytic core of the γ-secretase complex [39]-responsible for APP cleavage and Aβ generation. Mutations in PS1 represent the most common genetic cause of AD[5, 40]. Here, we demonstrate that PS1 interacts with mitochondrial proteins and regulates mitochondrial functions independently of its role in γ-secretase activity. Mitochondrial dysfunction is a well-established feature of AD, characterized by impaired energy metabolism, disrupted calcium homeostasis, altered mitochondrial dynamics, and ROS accumulation[15, 18, 19, 41]. Aβ and its precursor protein APP have been shown to disrupt mitochondrial energy metabolism and dynamics. For instance, pathogenic Aβ_42_ accumulates in mitochondria and impairs mitochondrial energy metabolism[42]. Neuronal mitochondria exhibit fragmented networks in AD, accompanied by defective mitophagy, leading to accumulation of damaged mitochondria and subsequent neuronal death[17, 43]. Additionally, mtDNA damage, which compromises bioenergetics and can trigger immune responses and apoptosis, is increasingly implicated in AD[18, 44, 45]. Recent studies have highlighted the importance of MICOS assembly in both physiological and pathological contexts.[46] We speculate that PS1 may facilitate proper MICOS organization, and its γ-secretase-independent functions also contributes to AD pathogenesis. Our results thus establish a novel link between PS1 and mitochondrial dysfunction, offering a novel perspective on AD progression.

Several limitations of this study should be noted. First, we focused on the loss of PS1 and its effects on mitochondrial structure and function. Since PS1 mutations are frequently identified in AD patients, future studies should examine how disease-associated PS1 mutant affect mitochondrial integrity and functions. Second, as both NTF and CTF fragments of PS1 were detected in mitochondria, their individual roles in regulating cristae architecture warrants further investigation. Additionally, the exact mechanism by which ATAD3A oligomerization impairs mitochondrial inner membrane integrity remains to be clarified.

## Supporting information

Supplemental realted to Fig5

## Acknowledgements

We thank Translational Medicine Core Facility of Shandong University for consultation and instrument availability (Zeiss LSM900 confocal microscope) that supported this work.

