## Supplemental realted to Fig5 for "Presenilin 1 (PS1) located at mitochondrial inner membrane regulates mitochondrial cristae junction proteins arrangement and cristae formation in HEK293 cells"

Supplementary Figure 1

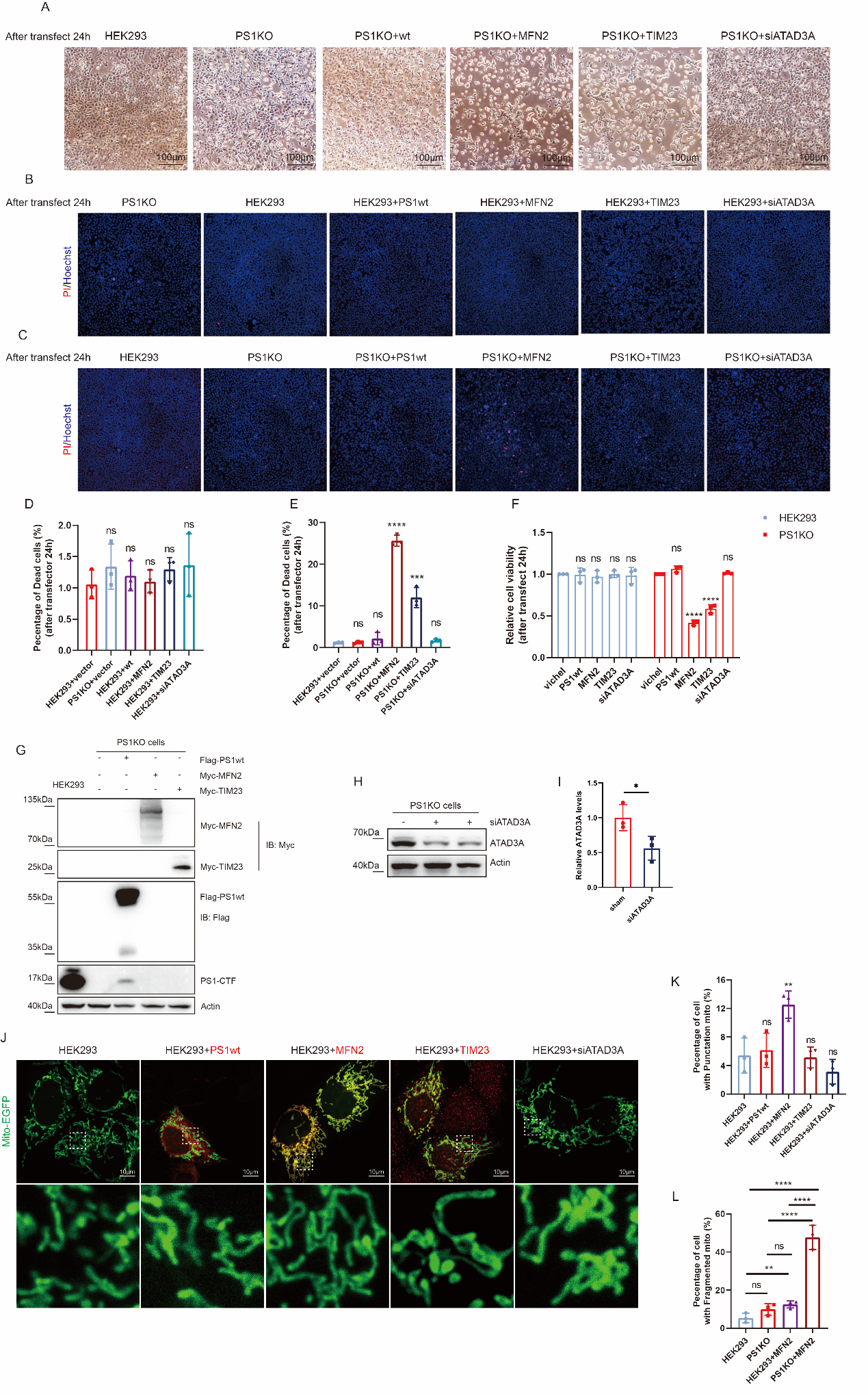

Fig. S1. (Related to Fig.5. Rescue attempts via MFN2/TIM23 overexpression or ATAD3A knockdown in PS1KO cells) (A) Bright‑field images of cells 24 h post‑transfection following seeding at equal density and overnight culture. (B, C) PI/Hoechst staining of HEK293 (B) or PS1KO (C) cells 24 h after overexpression of MFN2/TIM23 or knockdown of ATAD3A. (D, E) Quantification of cell death rates from (B) and (C), respectively. (F) Cell viability measured by CCK‑8 assay 24 h post‑transfection. (G) Western blot analysis of Myc‑MFN2 and Myc‑TIM23 overexpression. (H) Western blot confirming ATAD3A siRNA knockdown efficiency. (I) Quantification of ATAD3A levels from (H). (J) Mitochondrial morphology visualized by Mito‑EGFP in HEK293 cells following overexpression of PS1wt, MFN2, or TIM23, or after ATAD3A knockdown. (K) Percentage of cells exhibiting punctate mitochondria (defined as completely fragmented organelles). (L) Summary of cells with punctate morphology in HEK293 and PS1KO groups (data integrated from Fig. 5K and this figure). Data are presented as mean ± SD of at least three independent experiments. ✱*p* <0.05. ✱✱p <0.01. ✱✱✱p <0.005. ✱✱✱✱*p* <0.001.n.s. not significant.
